# An important role of the interplay between *Bdnf* transcription and histone acetylation in epileptogenesis

**DOI:** 10.1101/2020.09.01.277327

**Authors:** Agnieszka Walczak, Iwona Czaban, Anna Skupien, Katarzyna K. Pels, Andrzej A. Szczepankiewicz, Katarzyna Krawczyk, Błażej Ruszczycki, Grzegorz M. Wilczynski, Joanna Dzwonek, Adriana Magalska

## Abstract

Brain-Derived Neurotrophic Factor is one of the most important trophic proteins in the brain. The role of this growth factor in neuronal plasticity, in health and disease, has been extensively studied. However, mechanisms of epigenetic regulation of *Bdnf* gene expression in epilepsy are still elusive. In our previous work, using a rat model of neuronal activation upon kainate-induced seizures, we observed a repositioning of *Bdnf* alleles from the nuclear periphery towards the nuclear center. This change of *Bdnf* intranuclear position was associated with transcriptional gene activity.

In the present study, using the same neuronal activation model, we analyzed the relation between the percentage of the *Bdnf* allele at the nuclear periphery and clinical and morphological traits of epilepsy. We observed that the decrease of the percentage of the *Bdnf* allele at the nuclear periphery correlates with stronger mossy fiber sprouting - an aberrant form of excitatory circuits formation. Moreover, using *in vitro* hippocampal cultures we showed that *Bdnf* repositioning is a consequence of the transcriptional activity. Inhibition of RNA polymerase II activity in primary cultured neurons with Actinomycin D completely blocked *Bdnf* gene transcription and repositioning observed after neuronal excitation. Interestingly, we observed that histone deacetylases inhibition with Trichostatin A induced a slight increase of *Bdnf* gene transcription and its repositioning even in the absence of neuronal excitation. Presented results provide novel insight into the role of BDNF in epileptogenesis. Moreover, they strengthen the statement that this particular gene is a good candidate to search for a new generation of antiepileptic therapies.

## Introduction

Brain-Derived Neurotrophic Factor (BDNF) is one of the most important neurotrophins in the brain. Acting *via* its synaptic receptor Tropomyosin-related kinase B (TrkB), BDNF is involved in neuronal differentiation, survival, and synaptic plasticity (1–3). Thus, BDNF plays an important role in the number of neurological and psychiatric disorders such as Parkinson’s disease (4), schizophrenia (5), depression (6), bipolar disease (7), and epilepsy (8–10).

Currently, it is known that BDNF is involved in the event of aberrant synaptic plasticity called mossy fiber sprouting, observed in temporal lobe epilepsy, which is one of the most common types of epilepsy in adults (11). The level of both BDNF protein (12) and mRNA (13–15) were described to be elevated after seizures in the temporal lobe and hippocampi of epileptic patients. Experiments performed on animals and *in vitro* models showed that BDNF causes hypertrophy of granule neurons (16) and increased mossy fiber branching (17). Moreover, intrahippocampal infusion of BDNF induced mild seizures with the development of mossy fiber sprouting (18). Those findings support the involvement of BDNF in the aberrant synapse formation in temporal lobe epilepsy, however, underlying molecular mechanisms are still not clear.

The current trend in neuroscience is to look for mechanisms of neuronal functioning at a gene expression level. BDNF encoding gene is a so-called delayed, immediate-early gene induced in a later phase of neuronal activation (19). It consists of 9 exons differentially expressed in humans, mice, and rats (20). Regulation of BDNF expression after neuronal excitation has been quite well understood at the level of transcription factors and chromatin modifications. It is known that 3’ of the protein-coding exon is spliced to one of the eight of 5’ untranslated exons. Each of the 5’ exons is controlled by its unique promoter (20–22). Moreover, *Bdnf* can be epigenetically down-regulated through DNA methylation and histone deacetylation. The aforementioned epigenetic changes result in the recruitment of REST/NRSF (23) and MeCP2 (24, 25) and chromatin remodeling. Conversely, the up-regulation of the gene can be triggered by DNA demethylation and/ or histone acetylation, which was already presented in both *in vitro* (20) and *in vivo* studies (26). Importantly, the level of histone H3 acetylation at the *Bdnf* promoters IV and VI may underlie sustained up-regulation of transcription following chronic electroconvulsive shock (27). Fukuchi and colleagues (28) showed that valproic acid, an inhibitor of histone deacetylases commonly used antiepileptic drug, increases expression of *Bdnf* under control of promoter I.

Studies in the last decade have shown that the genome is spatially organized within the nucleus (29). Rearrangements of chromatin are involved in the regulation of gene expression, and the radial position of genes reflects their expression (30). In differentiated cells, the nuclear periphery is a repressive environment, where heterochromatin is recruited to the nuclear lamina (31). Artificial localization of gene at the nuclear periphery, by tethering to the inner nuclear membrane, is sufficient to induce silencing of its expression (32). The role of chromatin structure in the regulation of gene expression in neurons remains still unexplored. Crepaldi and colleagues (33) showed that activity-dependent genes, including *Bdnf*, were repositioned to transcription factories after KCl induced depolarization in cultured cortical neurons. Moreover, in our previous studies (34) we showed that during neuronal excitation and epileptogenesis *Bdnf* alleles had been detached from the nuclear lamina and repositioned from the nuclear periphery toward the nuclear center. The observed phenomenon was associated with changes in *Bdnf* expression. However, it was not clear whether *Bdnf* repositioning is a cause or a consequence of the transcriptional activity of the gene. Therefore, in the current study, we are addressing this interesting question.

## Materials and methods

### Animals

The experiments were performed on young, adult male Wistar rats, weight 170–250 g, obtained from Mossakowski Medical Research Centre, Polish Academy of Sciences. Animals were kept under a 12 h light/dark cycle, with unlimited food and water supplies. All procedures were performed with the consent of the 1st Local Ethical Committee in Warsaw (Permission number LKE 306/2017).

### Induction of seizures

Seizures were evoked by three doses of kainate (5 mg/kg, Sigma-Aldrich) (0.5% solution in saline, pH 7), administered intraperitoneally in 1 h intervals, and scored as described by Hellier et al (35). The animals were taken for further studies regardless of whether they fulfilled the criterion of the full status epilepticus, or not (35). To reduce kainate-induced mortality, diazepam (25mg/kg) was administrated intraperitoneally 3-6 hours after seizures onset.

### Estimation of clinical and morphological traits

Clinical and morphological traits such as the intensity of seizures, ruffling of the fur, forelimb clonus, body, and head tremor, loosing of posture, immobility and aggression have been estimated for 6 hours after kainate injection, and for 4 hours/day in subsequent 4 weeks. The intensity of seizures was scored according to the 6-grade modified Racine’s scale (0-lack of seizures to 5-fully developed tonic-clonic seizures with loss of posture) (35). Ruffling of the fur was scored in a 3-grade scale (0-lack of ruffling to 2-extensively ruffled fur). Forelimb clonus was scored according to a 7-grade scale (0-lack of clonus to 6-very strong forelimb clonus). Body and head tremor was scored according to 6-grade scale (0-lack of tremor to 5-very strong tremor). Loss of posture was scored according to the 4-grade scale (0-lack of loss of posture to 4 constant loss of posture). Immobility was scored according to the 10-grade scale (0-completely active to 9-lack of activity). Aggression was scored on a 4-grade scale (0-lack of aggression to 3-extreme aggression). To minimize the bias, the scoring of clinical parameters was performed by two independent observers. Mossy fiber sprouting was verified by immunofluorescent staining for synaptoporin and scored in 5-grade scale (0-lack of sprouting to 4-very strong sprouting).

### Primary neuronal hippocampal cultures and treatment

Primary neuronal cultures were prepared from the hippocampi of P0 rat brains as described previously (36). Chemical long term potentiation (cLTP) was initiated by stimulating the cells for 2 hours with 50 μM of picrotoxin, 50 μM forskolin, and 0.1 μM rolipram (all from Sigma-Aldrich). Transcription was inhibited by 2h exposure to 8mg/ml of Actinomycin D (Sigma-Aldrich). Histone deacetylases were inhibited by 12h treatment with 200nM Trichotstatin A (Abcam).

### Fluorescent *in situ* hybridization

Fluorescent *in situ* hybridization was performed according to the protocol of Cremer et al. (2008) (37) on 30 μm-thick brain cryosections of the 4% paraformaldehyde-perfused, 8 kainate-treated and 4 control animals, as well as on 3 independent primary hippocampal neuronal cultures fixed with 4% paraformaldehyde. As templates for *Bdnf* FISH probes, CH230-449H21 BAC obtained from Children’s Hospital Oakland Research Institute were used. Probes were verified on rat metaphase spreads. The probes were labeled using the standard nick-translation procedure. Biotinylated probes were detected using Alexa Fluor 488-conjugated avidin (Invitrogen), followed by FITC-conjugated rabbit anti-avidin antibody (Sigma-Aldrich).

### Immunostaining

Immunostaining for synaptoporin was performed on 30 μm-thick brain cryosections of the 4% paraformaldehyde-perfused animal using standard immunofluorescent staining protocol (38). 1 μg/ml of rabbit polyclonal anti synaptoporin (Synaptic Systems) antibody was used. The intensity of immunostaining was estimated using the 5-grade scale (0-no staining to 4-strong staining). The neuronal damage was examined by staining with Fluoro-Jade B (Millipore) according to the method of Schmued et al (39).

### Image acquisition

Fluorescent specimens were examined under TCS SP8 confocal microscope (Leica) or Zeiss 800 confocal microscope (Zeiss), by sequential scanning of images, with a pixel size of 80 nm and axial spacing of 210 nm, using a PlanApo oil-immersion 63 (1.4 numerical aperture) objective.

### Quantitative image analysis

The minimal distance from the nuclear periphery and *Bdnf* alleles in neuronal nuclei of animals was calculated using custom-written software, Segmentation Magick, described in Ruszczycki et al. (40). For neuronal cultures, custom-written software Partseg was used, described in (41). At least 140 nuclei, from 3 independent experiments were analyzed for each experimental variant.

### Real-time reverse transcriptase-PCR for Bdnf mRNA

Total cellular RNA was isolated from three independent hippocampal primary cultures using RNeasy Mini Kit (Qiagen) according to the manufacturer’s procedure. 1 μg of RNA was subjected to RT reaction using the SuperScript first-strand synthesis system for RT-PCR (Invitrogen) according to the manufacturer’s protocol. PCR was performed using SYBR Green PCR Master Mix (Thermo Fischer Scientific). Forward and reverse primers, were respectively: 5-CCATAAGGACGCGGACTTGTAC and 5-AGACATGTTTGCGGCATCCAGG.

### Statistical analysis

The data were obtained from 3 independent batches of neurons and analyzed using Welch’s Anova test. The differences in the percentage of the gene and the nuclear periphery were analyzed using chi-square (all 4 groups together) and Fisher’s exact test for post-hoc pairwise comparison. The details are given in the figures’ legends. The statistical analysis was performed using GraphPad Prism software.

## Results

### Morphological traits of the kainate model of temporal lobe epilepsy and clinical implications of *Bdnf* repositioning in hippocampal granule neurons

To verify the kainate model of temporal lobe epilepsy in rats, animals were carefully observed for 6 hours after a kainate injection, and 4 hours/day in the subsequent 4 weeks. The intensity of seizures has been assessed according to modified 6-grade Racine’s scale (0-lack of seizures-5-fully developed tonic-clonic seizures with a loss of posture). Additionally, clinical traits such as the intensity of seizures, fur ruffling, forelimb tonus, body and head tremor, loss of posture, immobility, and aggression have been rated. Moreover, hippocampal specimens from the aforementioned animals were analyzed for neuronal damage in DG granule cell layer and CA3 pyramidal layer by Fluoro-jade B staining and for mossy fiber sprouting in DG molecular layer by synaptoporin staining.

More than 60% of animals underwent *status epilepticus* (fully developed tonic-clonic seizures with a loss of posture, Fig. 1A, Table 1), and the remaining animals showed moderate seizure symptoms like head tremors (wet dog shaking) and fur ruffling. In 4 weeks following administration of kainate 4 out of 8 animals showed fur ruffling, 4 out of 8 demonstrated forelimb tonus, all animals exhibited body and head tremor, 4 out of 8 showed loss of posture, 6 out of 8 - immobility and 4 out of 8 - aggression (for details see Table 1).

**Figure 1.**
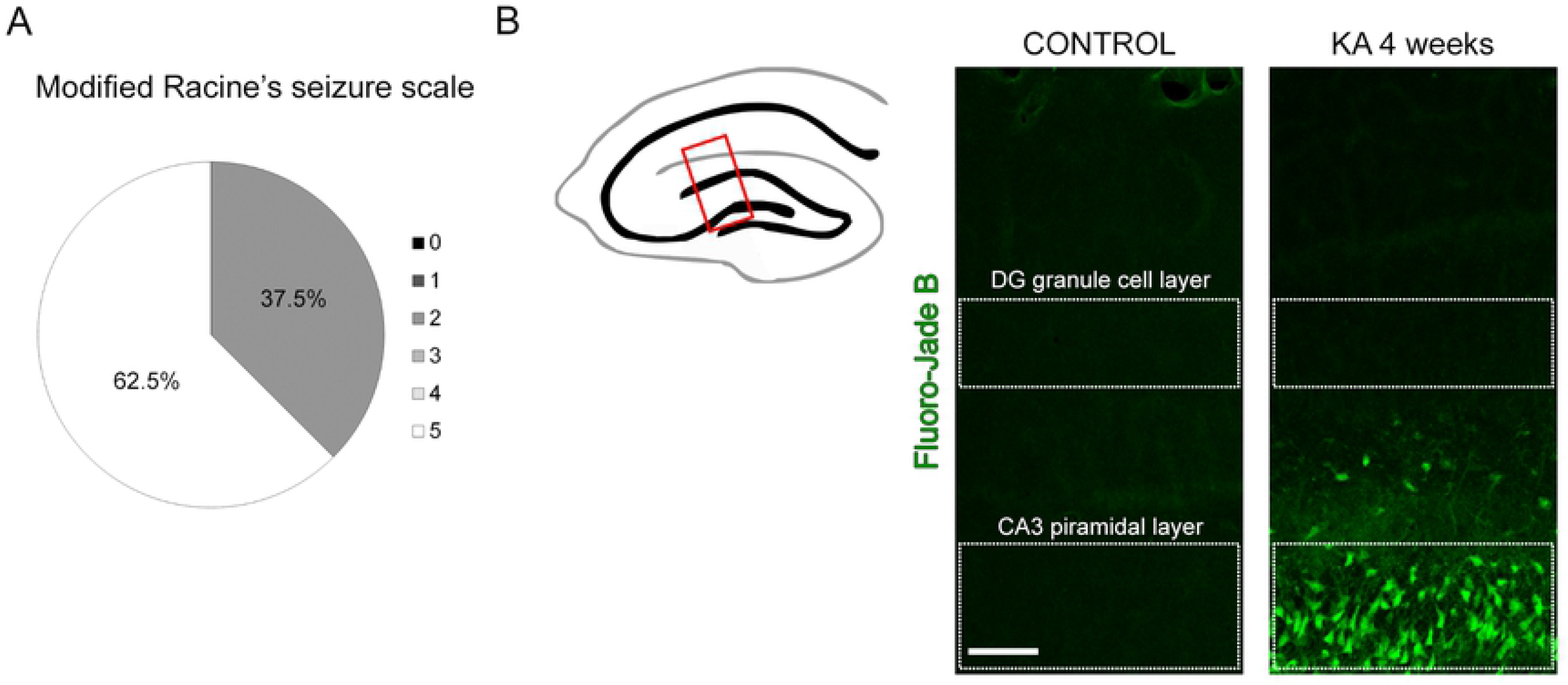
Kainate induced seizures and neuronal damage. A) Percentage of animals showing seizures after kainite administration. The intensity of seizure was scored according to the 6-grade modified Racine’s scale (0-lack of seizures - 5-fully developed tonic-clonic seizures with loss of posture). B) Fluoro-jade B staining (depicted in green) showing neuronal damage in CA3 region, but not DG region of the hippocampus of kainite treated animals. Scale Bar: 100 μm

**Table 1.**
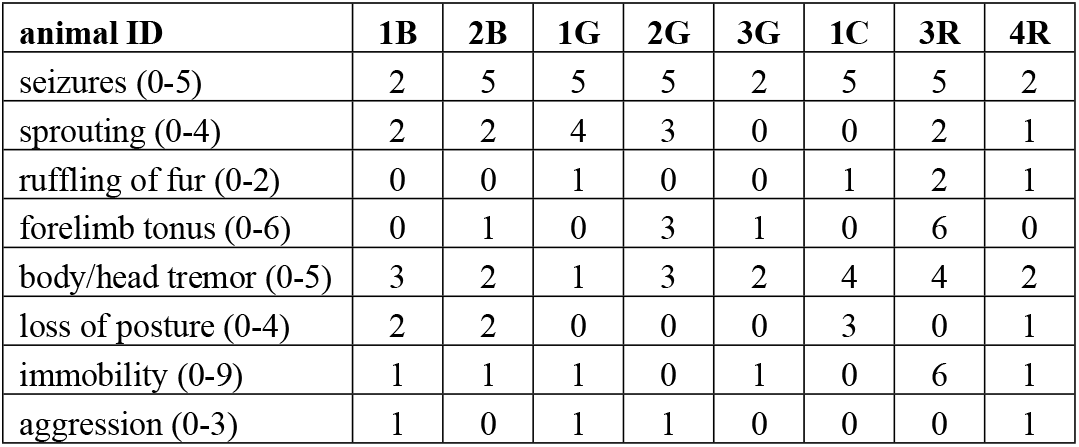
Estimation of clinical and morphological traits of the kainate model of temporal lobe epilepsy. For all traits, scales are shown in brackets (see Materials and Methods for full details).

| animal ID | 1B | 2B | 1G | 2G | 3G | 1C | 3R | 4R |
| --- | --- | --- | --- | --- | --- | --- | --- | --- |
| seizures (0-5) | 2 | 5 | 5 | 5 | 2 | 5 | 5 | 2 |
| sprouting (0-4) | 2 | 2 | 4 | 3 | 0 | 0 | 2 | 1 |
| ruffling of fur (0-2) | 0 | 0 | 1 | 0 | 0 | 1 | 2 | 1 |
| forelimb tonus (0-6) | 0 | 1 | 0 | 3 | 1 | 0 | 6 | 0 |
| body/head tremor (0-5) | 3 | 2 | 1 | 3 | 2 | 4 | 4 | 2 |
| loss of posture (0-4) | 2 | 2 | 0 | 0 | 0 | 3 | 0 | 1 |
| immobility (0-9) | 1 | 1 | 1 | 0 | 1 | 0 | 6 | 1 |
| aggression (0-3) | 1 | 0 | 1 | 1 | 0 | 0 | 0 | 1 |

Fluoro-jade B staining showed no neuronal damage in DG granule cell layer and extensive cell death in CA3 pyramidal layer in all animals (Fig. 1B). Extend of mossy fiber sprouting was estimated by the intensity of synaptoporin immunostaining (Fig. 2A and B). In 6 out of 8 animals, it was possible to distinguish mossy fibers stained against synaptoporin (scored 1 - weak sprouting to 4 - very strong sprouting, in Fig. 2A, B, and Table 1), where 25% showed moderate to very strong sprouting (scored ≥3). The level of sprouting was independent of the intensity of the initial seizures (right after kainate injection). All tested animals showed behavioral and morphological traits of epilepsy within 4 weeks from kainite treatment.

**Figure 2.**
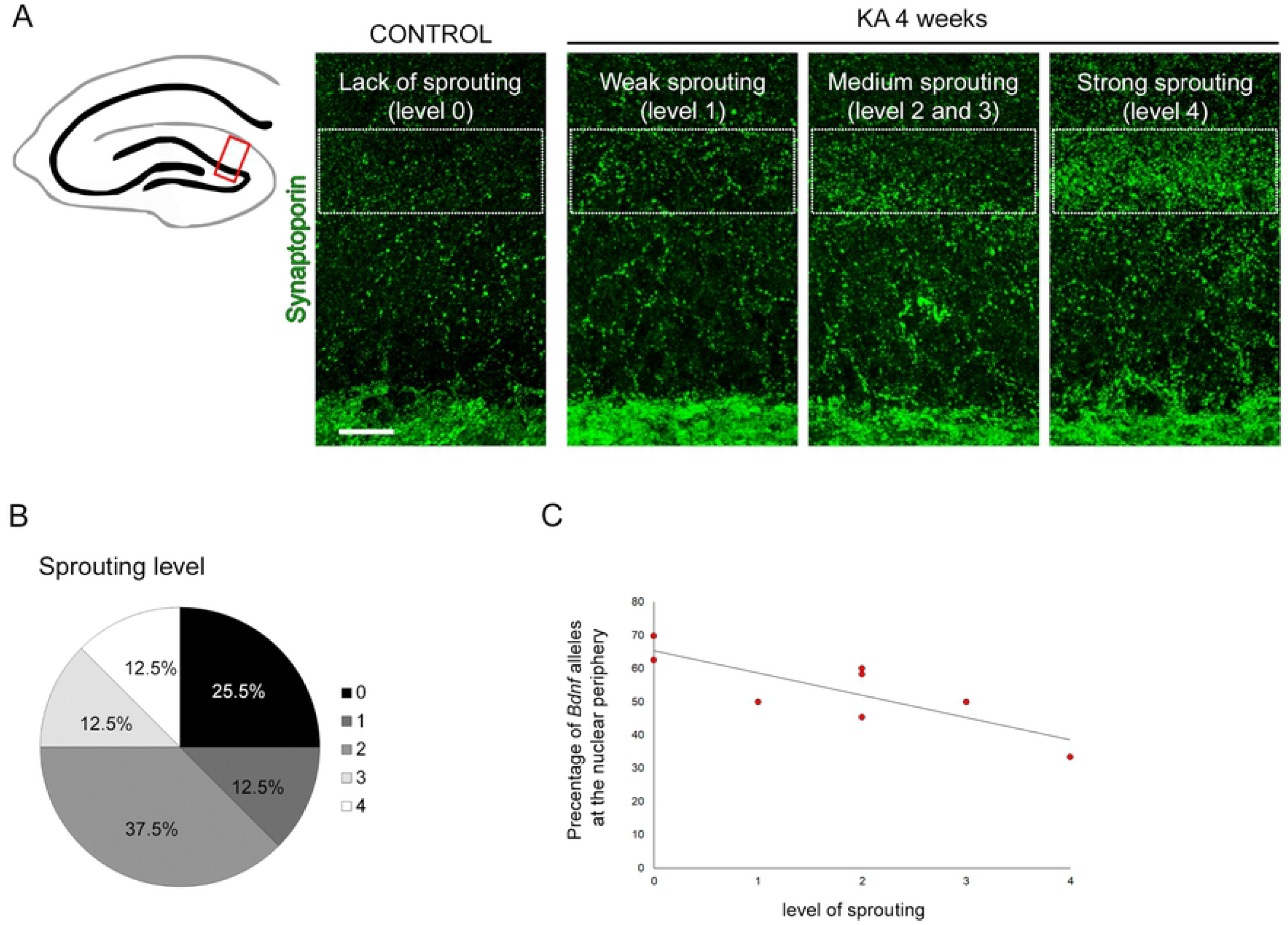
Correlation between the intensity of sprouting and the percentage of the nuclei with *Bdnf* alleles at the nuclear periphery. A) Mossy fiber sprouting was verified by immunofluorescent staining for synaptoporin in the molecular layer of DG region of the hippocampus. Representative pictures of different levels of sprouting of control animals and 4 weeks after administration of kainate are shown. B) Percentage of animals showing different levels of synaptoporin staining intensity scored in 5-grade scale (0-lack of sprouting-4 very strong sprouting). C) Correlation between the level of sprouting in DG 4 weeks from the administration of kainate, measured in 5-grade scale, and percentage of BDNF alleles localized at the nuclear periphery. Scale Bar: 100 μm.

In our previous work (34) using the same model of neuronal activation upon kainate-induced seizures, we have observed repositioning of the *Bdnf* alleles in hippocampal granule neurons. We have shown that transcriptionally inactive *Bdnf* is attached to a nuclear lamina and localized at the nuclear periphery, after neuronal activation it has repositioned towards the nuclear center. Here we observed, that the level of sprouting correlated (R = −0.81, Pearson correlation) with the percentage of the nuclei with *Bdnf* allele localized at the nuclear periphery (Fig. 2 C). In animals with the strongest level of sprouting, less than 50% of nuclei had *Bdnf* alleles present at the nuclear rim, while lack of sprouting correlated with the higher percentage of nuclei showing *Bdnf* localized in proximity to the nuclear envelope.

### The causal relationship between *Bdnf* allele transcriptional activity and repositioning

To further investigate a cause of the intranuclear reposition of *Bdnf* and its relationship with BDNF transcription, we used an *in vitro* model of neuronal excitation based on hippocampal dissociated cultures and a chemical model of long-term potentiation (cLTP) (42–44). cLTP evoked by picrotoxin, forskolin, and rolipram was proven to be non-toxic for neurons and to induce a program of gene expression similar to the one observed in a brain upon stimulation. We observed that 2 hours after the initiation of long term potentiation, *Bdnf* expression was significantly (4 times) increased compared to the control (Fig. 3A). At the same time, the *Bdnf* allele had repositioned toward the center of a cell nucleus (Fig. 3B-D). In the control cells (treated with a solvent alone) the *Bdnf* alleles were most frequently positioned at a nuclear margin, with 76.4 % of alleles located 350 nm or less from a nuclear border (Fig. 3 C and D, blue bars). This distance is an approximate microscope resolution limit, hence it has been chosen as an indicator of allele proximity to the nuclear border, as previously described (34). In neurons activated with cLTP, we observed a distinct repositioning of the *Bdnf* gene from the nucleus periphery towards the nucleus center (Fig. 3C and D, orange bars). The percentage of *Bdnf* alleles localized closer than 350 nm to the nuclear margin was significantly lower than in the control group (62.9 % Fisher’s exact tests). To verify whether *Bdnf* allele repositioning is a cause or a consequence of transcriptional activity, we performed experiments with the use of Actinomycin D, a potent inhibitor of RNA polymerase II. Preincubation with Actinomycin D for 2 hours was sufficient to inhibit *Bdnf* expression upon stimulation with cLTP (6 fold decrease compared to cLTP treatment, Fig 3A). Inhibition of transcription completely blocked *Bdnf* repositioning upon cLTP treatment (Fig 3B-D). Percentage of *Bdnf* alleles located at the nuclear periphery upon Actinomycin D and cLTP treatment was similar to control cells. Presented results show that transcriptional activity is a cause of *Bdnf* repositioning.

**Figure 3.**
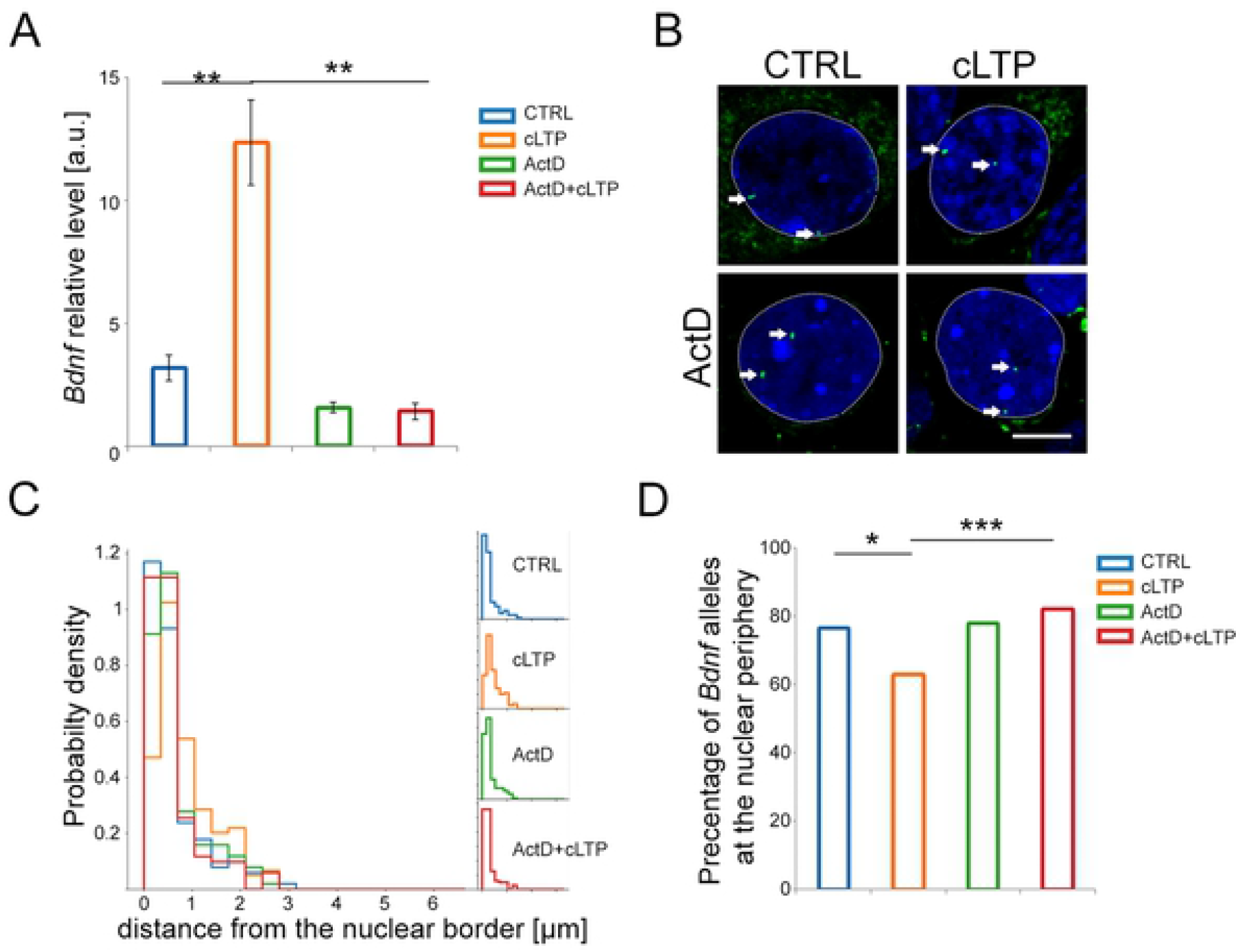
The causal relationship between *Bdnf* allele transcriptional activity and repositioning. (A) The graph shows the expression of *Bdnf* relative to *Gapdh* in the hippocampal neurons incubated for 2 hours with DMSO vehicle (CTRL, blue bar) or cLTP (orange bar), incubated for 2 hours with Actinomycin D and 2 hours with DMSO (ActD, green bar) or cLTP (ActD+cLTP, red bar). Welch’s ANOVA test: ** p<0,01; error bars indicate standard error of the mean for 3 independent experiments (B) Representative picture of the nuclei of hippocampal neurons treated as described above. Hoechst 3342 staining for chromatin is shown in blue and FISH signals for *Bdnf* gene are shown in green (white arrows), margins of the nucleus are shown by a white, dotted line. Scale bar 5μm. (C) Quantitative analysis of the intracellular positions of *Bdnf* alleles in the nuclei of hippocampal neurons treated and color-coded as in A. The minimal distance between the respective alleles and nucleus surface is presented in the normalized histogram. (D) Percentages of *Bdnf* alleles localized < 350 nm to the nuclear surface are shown (Chi-square test, all groups, p<0.001; Fisher’s exact tests * p<0.05, *** p<0.001).

### Histone deacetylases are necessary for the attachment of *Bdnf* alleles to the nuclear lamina

Covalent modifications of chromatin were shown to be involved in the regulation of *Bdnf* gene expression (27, 45, 46). Data indicate, that histone deacetylases (HDAC) play an important role in this process (47, 48). Therefore, we have raised a question of HDAC involvement in *Bdnf* repositioning observed after neuronal stimulation. Preincubation of hippocampal cultures with Trichostatin A, which is commonly used HDAC inhibitor, slightly increased *Bdnf* transcription (Fig 4A) and was sufficient to induce *Bdnf* allele repositioning toward the nuclear center (Fig 4B-D). The percentage of *Bdnf* alleles localized at the nuclear periphery was significantly lower upon TSA treatment compared to the control group. Induction of cLTP after Trichostatin A treatment did not decrease the percentage of alleles located at the nuclear periphery, in comparison to cLTP or TSA alone. This result together with our previous observations (34) shows, that HDACs activity plays an important role in the attachment of *Bdnf* alleles to the nuclear periphery.

**Figure 4.**
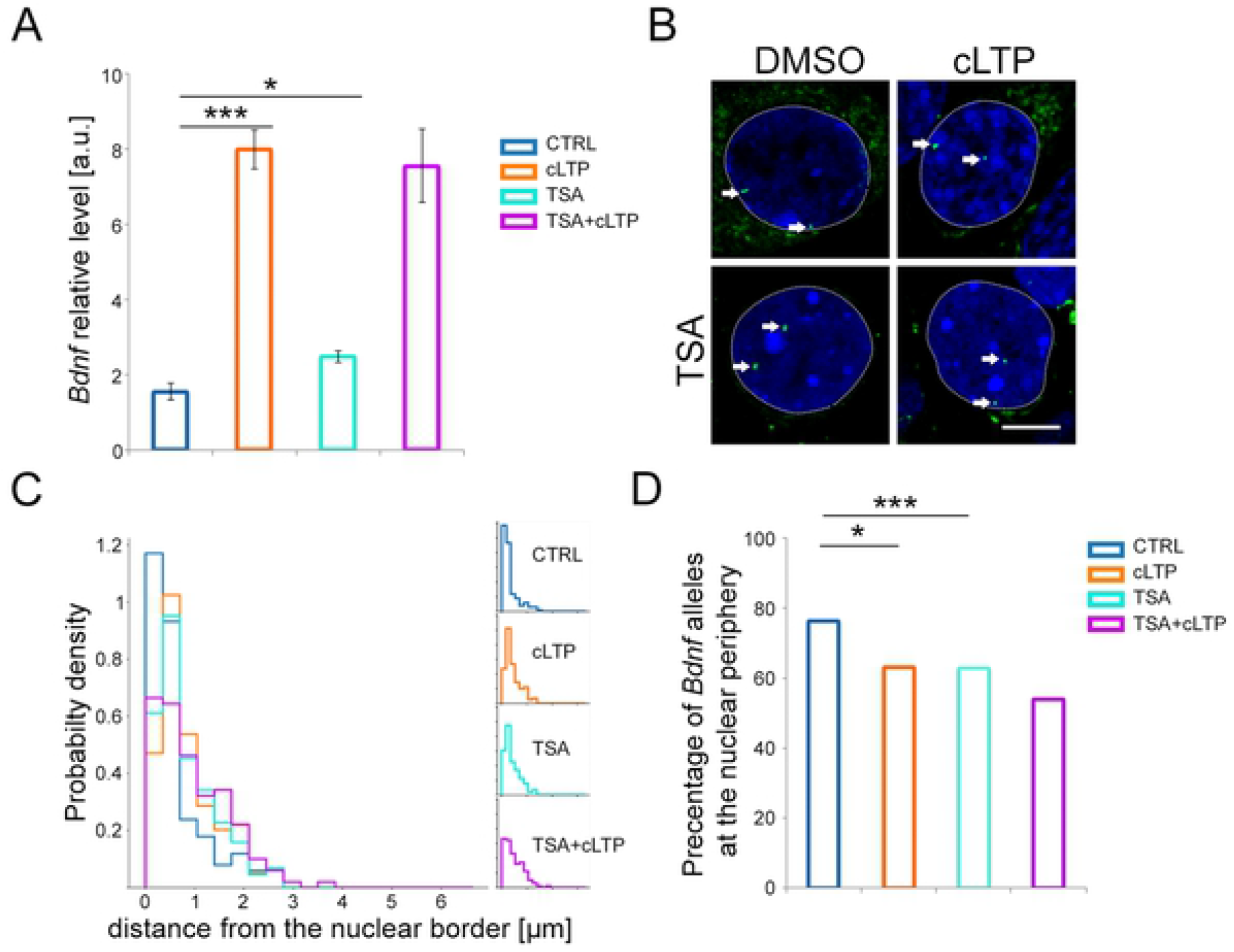
Importance of histone deacetylases in *Bdnf* repositioning. (A) The graph shows the expression of *Bdnf* relative to *Gapdh* in the hippocampal neurons incubated for 2 hours with DMSO vehicle (blue bar) or cLTP (orange bar), incubated for 2 hours with TSA and 2 hours with DMSO (TSA, cyan bar) or cLTP (TSA+cLTP, magenta bar). Welch’s ANOVA test: * p<0,05, *** p<0.001; error bars indicate standard error of the mean for 5 independent experiments (B) Representative picture of the nuclei of hippocampal neurons treated as described above. Hoechst 3342 staining for chromatin is shown in blue and FISH signals for *Bdnf* gene are shown in green (white arrows), margins of the nucleus are shown by a white, dotted line. Scale bar 5μm. (C) Quantitative analysis of the intracellular positions of *Bdnf* alleles in the nuclei of hippocampal neurons treated and color-coded as in A. The minimal distance between the respective alleles and nucleus surface is presented in the normalized histogram. (D) Percentages of nuclei with the minimum distance between the respective alleles and nucleus surface < 350 nm are shown (Chi-square test, all groups, p<0.001; Fisher’s exact tests * p<0.05, *** p<0.001).

## Discussion

Three-dimensional organization of the chromatin in the cell nucleus is a higher-order regulator of gene expression (49–51). The role of the nuclear lamina as a transcriptional activity regulating the compartment is already quite well explored (52–54). However, the knowledge regarding the particular mechanisms that are responsible for driving detachment of the genes from the nuclear lamina is still underinvestigated (55).

To study a relationship between *Bdnf* repositioning and morphological and clinical epileptic traits, we used an animal model of temporal lobe epilepsy. In particular, we used a kainate model of epilepsy because it resembles morphological traits of temporal lobe epilepsy such as mossy fiber sprouting in dentate gyrus and neurodegeneration in CA1 and 3 regions of the hippocampus (56, 57). To study mechanisms of *Bdnf* allele repositioning we decided for *in vitro* model of neuronal stimulation. *In vitro* model was chosen to avoid problems with the distribution of the Actinomycin D (58) through the blood-brain barrier and decrease the number of sacrificed animals. Transcriptomic studies by Dabrowski and collaborators showed that the expression pattern in excited neurons *in vivo* is sustained in *in vitro* model (59). Moreover, Szepesi and collaborators (2013) (42) showed that induction of excitation by rolipram, forskolin, and picrotoxin induced the formation of dendritic spine protrusions, which resembled *in vivo* formation of new synapses.

The nuclear lamina is known as a transcriptionally repressive nuclear compartment (60). Williams and colleagues (2007) (61) and Peric-Hupces and colleagues (2010) (62) presented rearrangement of the interactions between chromatin and nuclear lamina and the association of such phenomenon with transcription during differentiation of embryonic stem cells into neurons. However, the number of reports of the relationship between the rearrangement of chromatin architecture and transcription in fully differentiated neurons is very limited. In a presented study we confirmed using *in vitro* model our previous finding (34) that after neuronal excitation *Bdnf* alleles are repositioned toward the nuclear center. However, in the used *in vitro* model, the percentage of alleles localized at the nuclear periphery was higher in control cells, and difference to stimulated neurons was smaller than presented in the aforementioned paper. Such discrepancy might be a result of a more uniform environment in cell culture compared to *in vivo* situation. Additionally, kainate treatment, used *in vivo* studies induces much stronger neuronal excitation than chemically induced LTP, that had been applied *in vitro*. In the presented study we showed that inhibition of transcription prevents *Bdnf* repositioning in neuronal nuclei after stimulation. It suggests that depolarization of the neuronal membrane is not sufficient for the detachment of *Bdnf* alleles, but the transcriptional activity itself is necessary for full repositioning. Our finding is consistent with expertise presented by Crepaldi and colleagues (2013), who showed that in cultured cortical neurons depolarization induced by KCl stimulation evoked repositioning of activity-induced genes, including *Bdnf* into transcription factories (33). Moreover, the study showed that repositioning is controlled by transcription factor TFIIIC. Our current finding complements the previous study (34), where we showed that *Bdnf* repositioning is associated with changes in transcription level during epileptogenesis. Both of those studies support the hypothesis that *Bdnf* allele repositioning acts as a kind of molecular memory of the cell to prepare neurons for future activation. The finding is also in agreement with the report by Ito and colleagues (2014) showing, that loss of three-dimensional architecture in neuronal nuclei leads to impaired transcription of several genes (63).

Most probably, the presented mechanism is not the only one involved in the *Bdnf* repositioning. Most likely several pathways must be orchestrated to activate *Bdnf* transcription and detachment of gene from the nuclear lamina, as well as its reposition toward the nuclear center. One possible mechanism may involve cohesins, which are well-known genome organizers (64) (and references therein). In Cornelia de Lange Syndrome which belongs to cohesinopathies and is associated with epileptic seizures (65, 66) acetylation of cohesins is impaired due to mutation in the gene encoding histone deacetylase HDAC8 (67). The cohesin-dependent mechanism may be at least in part responsible for *Bdnf* transcriptional activation, detachment, and repositioning, as CTCF, which acts in concert with cohesins (68) was shown to regulate the transcription of *Bdnf* (69). Also, transcription factors such as Serum Response Factor (SRF) seem to be good candidates responsible for *Bdnf* repositioning. It is known that SRF can regulate *Bdnf* transcription (70) and its deletion leads to increased epileptogenesis and decreased expression of activity-induced genes including *Bdnf* (71).

Furthermore, it was shown, that histone acetylation at the promoter region is necessary for activation of gene expression (72). Histone deacetylases are necessary to reverse this process and are taking part in creating a repressive zone by the nuclear lamina (73). In a presented study we showed, that inhibition of histone deacetylases with TSA leads to *Bdnf* transcription and repositioning. Similar activation of BDNF gene transcription upon TSA treatment was shown for Hek293 cell line (74) and HDACs involvement in *Bdnf* expression is well established (28, 47, 75–78). However, the exact mechanisms of HDACs participation in the higher-order mechanisms of transcriptional regulation of BDNF are unknown. One possible mechanism might involve Methyl-CpG-Binding protein 2 (MeCP2), which was also shown to be involved in BDNF expression. Activation of neurons leads to the decrease in CpG methylation of *the Bdnf* promoter region and dissociation of MeCP2 protein together with mSin3A silencing complex and histone deacetylase HDAC1 (25). Besides, MeCP2 binds to the inner nuclear membrane lamin B receptor (79) and Trichostatin A was shown to decrease MeCP2 expression (80), which could explain the association of transcriptionally inactive *Bdnf* to the nuclear lamina and its detachment upon TSA treatment. Mutations of the MeCP2 encoding gene are responsible for most cases of Rett Syndrome, a neurodevelopmental disorder, in which patients develop seizures (81). Additionally, the process of *Bdnf* repositioning might involve actin-based molecular motors, since studies by Serebryannyy and colleagues (2016) (82) reported that actin regulates the function of HDAC1 and HDAC2 and also can be involved in gene expression by association with RNA polymerase II.

Finally, in a presented study we attempted to investigate the involvement *Bdnf* repositioning in the pathogenesis of TLE in rats. We observed that the percentage of nuclei with the *Bdnf* allele at the nuclear periphery is highly correlated with the intensity of mossy fiber sprouting. That is consistent with an idea that BDNF takes part in sprouting events in epilepsy (17, 18). Moreover, in the presented study we used Actinomycin D which has been already in use in treatments of several types of cancer (83–85). However, in terms of the potential use of Actinomycin D for the treatment of brain diseases including epilepsy, its administration would be challenging due to difficulties of its distribution through the brain-blood barrier (58). Taking into account all the results together, the presented study supports the idea of Simonato (86), that *Bdnf* can be a very good target for novel anti-epileptic therapies. Additionally, our data support the suggestion of Jagirdar and colleagues (2015), that HDACs activity might be a good target for the treatment of TLE (87).

Conclusively, the presented results are consistent with a current trend in research on the pathogenesis of epilepsy and show that neuronal cell nuclei are an interesting target to search for mechanisms of epileptogenesis.

## Acknowledgments

JD was supported by the Polish National Science Centre grant No 2015/17/B/NZ4/02540, AS was supported by the Polish National Science Centre grant No 2018/29/B/NZ4/01473, AM was supported by the Polish National Science Centre grant No UMO-2015/18/E/ NZ3/00730. AAS was supported by the Polish National Science Centre grant No 2014/15/N/NZ3/04468. KKP was partially supported by the ETIUDA grant from the Polish National Science Centre No. UMO-2019/32/T/NZ4/00502.

